# The developmental transcriptome of contrasting Arctic charr (*Salvelinus alpinus*) morphs

**DOI:** 10.1101/011361

**Authors:** Jónes Gudbrandsson, Ehsan P. Ahi, Sigrídur R. Franzdottir, Kalina. H. Kapralova, Bjarni K. Kristjóson, Sophie S. Steinhaeuser, Ísak M. Jónesson, Valerie H. Maier, Sigurdur S. Snorrason, Zophonías O. Jónsson, Arnar Pálsson

## Abstract

Species showing repeated evolution of similar traits can help illuminate the molecular and developmental basis of diverging traits and specific adaptations. Following the last glacial period, dwarfism and specialized bottom feeding morphology evolved rapidly in several landlocked Arctic charr (*Salvelinus alpinus*) populations in Iceland. To initiate study of the genetic divergence between small benthic morphs and larger morphs with limnetic morphotype, we conducted an RNA-seq transcriptome analysis of developing charr. We sequenced mRNA from whole embryos at four stages in early development of two stocks with contrasting morphologies, the small benthic (SB) charr from Lake Thingvallavatn and Holar aquaculture (AC) charr.

The data reveal significant differences in expression of several biological pathways during charr development. There was also an expression difference between SB- and AC-charr in genes involved in energy metabolism and blood coagulation genes. We confirmed expression difference of five genes in whole embryos with qPCR, including *lysozyme* and *natterin like* which was previously identified as a fish-toxin of a lectin family that may be a putative immunopeptide. We also verified differential expression of 7 genes in developing heads, that associated consistently with benthic v.s. limnetic craniofacial morphology (studied in 4 morphs total). Comparison of Single nucleotide polymorphism (SNP) frequencies reveals extensive genetic differentiation between the SB- and AC-charr (60 fixed SNPs and around 1300 differing by more than 50% in frequency). In SB-charr the high frequency derived SNPs are in genes related to translation and oxidative processes. Curiously, three derived alleles in the otherwise conserved 12s and 16s mitochondrial ribosomal RNA genes are found in benthic charr.

The data implicate multiple genes and molecular pathways in divergence of small benthic charr and/or the response of aquaculture charr to domestication. Functional, genetic and population genetic studies on more freshwater and anadromous populations are needed to confirm the specific loci and mutations relating to specific ecological or domestication traits in Arctic charr.

## Introduction

Historical contingencies and chance shape organism during evolution [1, 2], but convergence in phenotype and molecular systems indicates that evolution is to some extent predictable [3, 4]. Identification of genes and variants that influence evolved differences is not a trivial task [5]. However by identifying genes and developmental pathways shaped by evolution, we can test models of repeatability or contingency [4]. An ideal system to study the role of chance and necessity in ecological evolution would be a species or species complex with readily observable phenotypic variation, living in tractable ecological setting, and most crucially showing parallel evolution of specific traits within/among species. Finches of the Galapagos islands and cichlids in the African great lakes are exciting multi-species systems in this respect [6–8] and threespine stickleback has emerged as a model “single species” system [9]. The diversity in feeding specialization of fishes provides great opportunities for studying adaptation and divergence at the developmental and genetic level.

### Transcriptomic analyses of fish biology and divergence

Multiple methods can be used to identify genes affecting adaptations [5, 15–17]. Recently transcriptomic methods have been used on many fish species to address evolutionary and ecological questions. For example microarrays were used to compare gene expression in anadromous and resident populations of brown trout (*Salmo trutta*), revealing that life history was a better predictor of gene expression in the liver than relatedness [18]. RNA-sequencing (RNA-seq) methods have been used to study non-model species such as the Mexican cavefish (*Astyanax mexicanus*), cod (*Gadus morhua*) brook charr (*Salvelinus fontinalis*) and Atlantic Salmon (*Salmo salar*) [19–24], addressing different questions (concerning evolution, molecular genetics, development and aquaculture). RNA-seq were applied to adult Canadian Arctic charr to study salinity tolerance, linking expression and quantitative trait loci [25]. Microarray studies of adult lake whitefish (*Coregonus cupleaformis*) pointed to parallel expression differences between benthic and limnetic forms [26]. Filteau *et al.* (2013, [27]) found that a set of coexpressed genes differentiated the two whitefish morphotypes, implicating Bone morphogenesis protein (BMP) signaling in the development of ecological differences in tropic morphology. An alternative approach in identifying pathways related to function or morphological differences is to study gene expression during development [28, 29].

Some northern freshwater fish species exhibit frequent parallelism in trophic structures and life history and in several cases are they found as distinct resource morphs [9–14]. One of these species, Arctic charr (*Salvelinus alpinus*), is well suited for studying the developmental underpinnings of trophic divergence and parallel evolution. Local adaptation has been extensively studied in the salmonid family, to which Arctic charr belongs [30]. The family is estimated to be between 63.2 and 58.1 million years old [31, 32]. Assembly and annotation of paralogous genes and multigene families is a major challenge. This task is made even more challenging because a whole genome duplication occurred before the radiation of the salmonid family [33–36] which has provided time for divergence of ohnologous genes (paralogous genes originated by whole genome duplication event). Furthermore, recent estimates from the rainbow trout (*Oncorhynchus mykiss*) genome suggest that ohnologous genes were lost at a rate of about 170 ohnologous genes per million years and by utilizing multiple data sources the genome assembly problem of this family can be solved [36]. *de novo* assembly of genomes and transcriptomes is complicated if many paralogs are present, such as in salmonids. Furthermore, for data with short reads, mapping to a related reference genome/transcriptome is recommended over de novo assembly [37].

### Molecular studies of the highly polymorphic Arctic charr

Following the end of the last graciai period, about 10.000 years ago, Arctic charr colonized northern freshwater systems [38]. It can be found as anadromous or lake/stream residents and exhibit high level of within species polymorphism [12, 38]. Resource polymorphism in charr correlates with ecological attributes [39–41]. For instance small charr with benthic morphology, are found in multiple lavaspring and pond habitats in Iceland [42], and a comparative study of Icelandic lakes [41] found that lakes with greater limnetic habitat, fewer nutrients, and greater potential for zooplankton consumption appeared to promote resource polymorphism. Some of the larger lakes contain two or more distinct morphs, typically limnetic and benthic forms. Multiple lines of evidence show that these differences stem both from environmental and genetic causes [43–47]. The best studied and most extreme example of sympatric charr morphs are the four morphs in Lake Thingvallavatn [48], two of which belong to a benthic morphotype, a large benthivorous (LB-charr) and a small benthivorous (SB-charr), and two limnetic morphs, a large piscivorous morph (PI-charr) and small planktivorous morph (PL-charr) [49]. PL- and PI-charr operate in open water and feed on free-swimming prey, planktonic crustaceans and small fish, respectively. The two benthic morphs mainly reside on the bottom, feeding mostly on benthic invertebrates. The SB-charr can utilize interstitial spaces and crevices in the littoral zone typically consisting of submerged lava, which due to its porous surface, offers a richer source of benthic invertebrate prey than do stones with smoother surfaces [50]. Several population genetics studies, using allozymes and mtDNA revealed no differences among charr populations [51–53], while studies on microsatellite markers and nuclear genes, reveled both subtle [54–56] and strong genetic differences among morphs [57]. Importantly Kapralova *et al.* (2011, [58]) concluded that small benthic morphs have evolved repeatedly in Iceland and that gene flow has been reduced between the PL and SB morphs in Lake Thingvallavatn since its formation approximately 10,000 years ago [59]. We also discovered genetic separation in immunological genes (*MHCIIα* and *cath2*) between morphs in Iceland and within the lake [57], consistent with ecologically driven evolution of immune functions. Recently it was shown that expression of mTOR pathway components in skeletal muscle correlates with the SB-charr form [60], but it is unknown whether genetic differentiation in those genes or at which stage of development they emerge. These studies relied on few genes, but as individual genes have distinct histories [61, 62], genome wide methods are needed to identify genes associated with divergence. Icelandic aquaculture charr (AC) was founded with fish from the north of Iceland, and has been bred at Holar University College since 1990 [63]. The Holar AC-charr has responded to artificial selection in growth and performance characteristics, and is now the dominant charr breed in aquaculture in Iceland. While clearly a derived form, it has retained general limnetic craniofacial morphotype (Figure 1). In this study we compare SB charr from Lake Thingvallavatn and AC charr because i) SB charr represents an extensively studied and derived form of charr, that has been separated from Anadromous fish for approx. 10,000 years, ii) of the availability of abundant AC material and iii) we wanted an extreme contrast, because of budget reasons we could only sequence 8 samples at the time. This rather extreme contrast is justified as the data and other studies ([64, 65]) building on this data illustrate (see discussion).

The aims of this project are threefold. First, to find genes and pathways related to the development of phenotypic differences between benthic and limnetic arctic charr morphs. Second, to screen for signals of genetic differentiation that may relate to divergence of benthic and limnetic charr. Third, we set out to verify a subset of the expression and genetic signals, in benthic and limnetic morphs. In this study we conduct RNA-sequencing using samples of developing offspring of two contrasting Arctic charr morphs, a small benthic charr from Lake Thingvallavatn and Icelandic aquaculture charr conforming to a limnetic morphotype. For each morph we sequenced RNA samples from embryos at three pre-hatching and one post-hatching stage. The data revealed significant expression differences between the two morphs, both involving specific loci and particular molecular systems. This enabled us also to identify candidate genetic changes and differential expression of developmental genes that may affect jaw and craniofacial traits which separate benthic and limnetic morphotypes in charr. The comparison of AC- and SB-charr thus uncovered genes and pathways that separate morphs within Lake Thingvallavatn. These results emphasize the broad utility of population level transcriptomics for studies aimed at dissecting the genetics and developmental aspects of morphological divergence.

**Figure 1. The two contrasting morphs used in this study.** Adult individuals of the two morphs; the Holar aquaculture charr above and the small benthic charr from Lake Thingvallavatn below. Differences in size, coloration and head morphology are apparent.

## Materials and Methods

### Sampling, rearing and developmental series

We set up crosses and reared embryos in the laboratory as previously described [64]. Embryos from four charr morphs were studied, an aquaculture charr (hereafter called AC-charr) from the Holar breeding program [63], and three natural morphs from Lake Thingvallvatn; small benthivorous (SB), large benthivorous (LB) and small planktivorous (PL) charr [66]. The first two, AC and SB-charr, which exhibit contrasting adult size and morphology (Figure 1), were collected in 2009 and material sent for developmental transcriptome sequencing. The latter two were sampled in 2010 and used for qPCR and SNP studies of selected genes. Briefly, in September 2009 we got material from spawning AC-charr from the Holar breeding program [63] and spawning SB-charr collected via gill netting in Olafsdrattur in Lake Thingvallavatn. Similarly in 2010 spawning SB-, LB- and PL-charr were collected from Lake Thingvallavatn. Fishing permissions were obtained from the Thingvellir National Park Commission and the owner of the Mjóanes farm. For each parent group eggs from several females were pooled and fertilized using milt from several males from the same group. Embryos were reared at ~5° C under constant water flow and in complete darkness at the Holar University College experimental facilities in Verid, Saudárkrókur. The water temperature was recorded twice daily and the average was used to estimate the relative age of the embryos using tau-somite (*τs*) [67]. Embryos and juveniles were sampled at designated time points, placed in RNAlater (Ambion) and frozen at –20°C. For the investigation of different tissues of adult aquaculture charr (AC) from Holar (fish size 20–25 cm) were used. Six randomly selected individuals were killed (cutting through spinal cord) and dissected and samples were taken from the skin, heart, liver, gills, spleen, intestine and kidney of each fish. The samples were placed in RNAlater (Ambion) and stored at –20°C. DNA for population genetic analyses was from our previous study [57].

Fishing in Lake Thingvallavatn was with permissions obtained both from the owner of the land in Mjoanes and from the Thingvellir National Park commission. Ethics committee approval is not needed for regular or scientific fishing in Iceland (The Icelandic law on Animal protection, Law 15/1994, last updated with Law 157/2012). Sampling was performed with Holar University College Aquaculture Research Station (HUC-ARC) personnel. HUC-ARC has an operational license according to Icelandic law on aquaculture (Law 71/2008), which includes clauses of best practices for animal care and experiments.

### RNA extraction and transcriptome sequencing

Embryos of AC- and SB-charr sampled in 2009 were used for transcriptome sequencing. For this we focused on the time covering development of pharyngeal arches and morphogenesis of the head, at 141, 163, 200 and 433 *τs* (post fertilization). For each combination of morphs and timepoints we pooled RNA from approximately 6 individuals. RNA extraction and following steps were performed as described earlier [64, 68]. The embryos were dechorionated and homogenized with a disposable Pellet Pestle Cordless Motor tissue grinder (Kimble Kontes, Vineland, NJ, USA) and RNA was extracted into two size-fractions using the Ambion mirVana kit (Life Technologies, Carlsbad, CA, USA). The high molecular weight fraction was further used for mRNA-seq and RNA quality was analysed using an Agilent 2100 Bioanalyzer (Agilent Technologies, Santa Clara, CA, USA). First and 2nd strand cDNA synthesis, fragmentation, adapter ligation and amplification were performed using the mRNA-Seq 8-Sample Prep Kit (Illumina, San Diego, CA, USA) according to manufacturer’s instructions. Sequencing was performed at DeCode genetics (Reykjavík, Iceland) using SOLEXA GAII technology (Illumina, San Diego, CA, USA).

The embryos sampled in 2010 were used for qPCR expression analyses. RNA was extracted from six whole embryos, in two replicates (2 X 3 fish) (AC and SB sampled at 161 and 200 *τs*). For the extraction of RNA from heads of AC, SB, LB and PL, 12 embryos (2 X 6) at 178, 200 and 216 *τs* were used. Embryos were dechorionated and decapitated in front of the pectoral fin. RNA extraction and cDNA preparation were performed as described previously [64]. Similarly RNA was extracted from a small piece (ca. 2 mm^2^) of skin, heart, liver, gill, spleen, intestine and liver from 6 adult AC-charr.

### Analyses of RNA-seq data and mapping to Salmon EST contigs

As no *S. alpinus* genome is available and de novo assembly of the 36 bp reads yielded excessive number of short contigs we chose to assess expression and genetic variation by mapping the reads to 59336 *S. salar* EST contigs from the SalmonDB [69, downloaded 22. March 2012] and the Arctic charr mitochondrial genome [61, NC_000861].

To estimate expression, reads were aligned with RSEM vs. 1.1.18 with default parameters. RSEM distributes reads that map to multiple locations to the most likely contig, using expectation maximization [70]. The read counts for contigs with the same annotation were pooled, because some genes were represented by more than one contig and due to whole genome duplication salmonids have closely related paralogous genes [34, 36]. Thus the expression tests are done on gene or paralog group level, instead of the contig level. In the remainder of the paper, gene will have this broader meaning, some genes are represented by one contig and others by two or more (indicated in all relevant tables). This brought the number of genes considered down to 16851. Lastly genes with fewer than 800 mapped reads in the entire dataset, were excluded from the analyses, yielding a total of 10496 genes.

A generalized linear model (GLM) with morph and developmental time as explanatory variables was used to find genes with different expression levels between the two charr morphotypes (groups) using the edgeR-package in R [71].

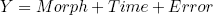

We could not test for interaction, as biological replicates were unavailable. To obtain further insight into the expression profiles of differently expressed genes, we preformed clustering analyses on log-transformed cpm-values (counts per million, cpm-function in edgeR). The values for each gene were scaled by mean and standard deviation and the euclidean distance used for the hclust-function in R [72] with the default settings. We used the Hypergeometric-test in goseq [73] to test for gene ontology enrichment. Since we pooled the read-count from different contigs we could not take gene length into account in those tests as would have been optimal.

The sequencing reads were deposited into the NCBI SRA archive under BioProject PRJNA239766 and with accession numbers: SRX761559, SRX761571, SRX761575, SRX761577, SRX761451, SRX761461, SRX761490 and SRX761501.

### Assessment of differential expression with qPCR

We previously identified suitable reference genes to study Arctic charr development [64]. Here we examined the expression of several genes in whole charr embryos, embryonic heads and adult tissues. Primers were designed using the Primer3 tool [74] and checked for self-annealing and heterodimers according to the MIQE guidelines [75] (S1 Table). Primers for genes with several paralogs were designed for regions conserved among paralogs. For natterin, primers for the different paralogs were designed to match regions differing in sequence. Relative expression was calculated using the 2^−∆∆*Ct*^ method [76]. For the calculation of relative expression of genes in whole embryos, the geometric mean expression of three reference genes, *β-Actin, elongation factor* 1*α* and *Ubiquitin-conjugating enzyme E2 L3*, was used for normalization. For visual comparisons among samples the normalized expression was presented as relative to the expression in AC at 161 *τs* (calibration sample). For the embryonic head samples *IF5A1* and *ACTB* were used as reference genes and a biological replicate of AC at 178 (*τs*) as the calibrator sample. Standard errors of relative expression were calculated from the standard errors (SE) of the ∆*C*_*T*_-values with the formula 2^−(∆∆*Ct*+*SE*)^ = minimum fold expression and 2^−(∆∆*Ct*−*SE*)^ = maximum fold expression. The statistical analysis was performed using the ∆*C*_*T*_-values with a two-way ANOVA with GLM function in R.

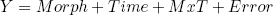

Normal distribution of residuals was confirmed for all data. For the study of expression in the embryonic head we followed a significant Morph effect in the ANOVA with Tukey’s post hoc honest significant difference test, on relative expression ratios (∆*C*_*T*_s).

### Polymorphisms in the Arctic charr transcriptome

For analysis of genetic variation we mapped the reads to the salmon contigs, this time using BWA [77] with a seed length of 25, allowing two mismatches. We re-mapped the reads since BWA allows short indels (RSEM does not), but disregarding them leads to many false SNPs close to indels. To extract candidate polymorphic sites from the Arctic charr transcriptome we ran VarScan2 [78] with minimum coverage of 50 reads and minimum minor allele frequency 0.1 on reads mapped to each *S. salar* contig for all of the 8 timepoints and morph combinations. This was done separately for reads that mapped uniquely to one contig only (UNI) and reads that mapped to two or more contigs (REP). These SNP-candidates were further processed in R [72]. SNP-candidates at 90% frequency or higher in all samples were disregarded, they reflect difference between Arctic charr and *S. salar* and are not the focus of this study. SNP-candidates with poor coverage in specific samples, that is coverage of 5 or fewer reads in 3 or 4 samples of each morph, were removed. As the SNP analysis was done on individual contigs, differences among paralogs appear in the data. But, since each sample is a pool of few individuals it is very unlikely that we have the same frequency of true SNPs in all the samples. This property was used to remove variants that are most likely due to expressed paralogs. Using Fisher exact tests to evaluate differences between samples only SNPs that were significantly different between samples with p-value lower than 0.05 (no multiple testing correction) were chosen for further examination. As equal cDNA input from individuals in sample cannot be assumed, due to expression differences among them and stochastic processes in sample preparation, read numbers were summed over the four samples for each morph for the comparison between the two groups. A conservative approach was taken to look for difference between morphs. We focused on SNP-candidates that showed difference in frequency between morphs without adjusting for multiple testing (Fisher exact test, *p* > 5%). We extracted the most interesting candidates by filtering on frequency difference between the morphs (delta). SNP-candidates with the highest frequency difference (delta > 95%) were manually processed and redundant candidates removed. A similar approach was used to mine for polymorphism in the Arctic charr mtDNA (NC_000861), using *S. salar* mtDNA as outgroup (NC_001960.1).

We wrote a python script to predict the impact of SNPs within the mRNA sequences. Polymorphisms where categorized according to their location (3’UTR, coding, 5’UTR), and those within the coding region into synonymous or non-synonymous.

### Verification of candidate SNPs

We chose 12 candidate SNPs for verification (see below). The candidates were verified using a similar approach as previously [57]. First we conducted genomic comparisons of the Salmon genome, ESTs and short contigs from the preliminary assembly of the Arctic charr transcriptome. This allowed us to infer the placement of the putative polymorphism in the locus, and design paralog specific primers for PCR (less than 1 kb amplicons) for verification of the 12 candidate SNPs. Each individual was genotyped by first amplifying the region of interest using PCR, followed by ExoSAP, direct sequencing (BigDye) and finally run on an Applied Biosystems 3500xL Genetic Analyzer (Hitachi). Raw data was base-called using the Sequencing Analysis Software v5.4 with KBTMBasecaller v1.41 (Applied Biosystems). Ab1 files were run through Phred and Phrap and imported to Consed for visual editing of ambiguous bases and putative polymorphisms and trimming primer. The fasta files were aligned with ClustalW online [79, http://www.ebi.ac.uk/Tools/msa/clustalw2/] and manually inspected in Genedoc [80]. All sequences where deposited to Genebank as Popsets under the accession numbers KP019972-KP020026.

### Comparative genomic analyses of sequence polymorphism

Comparative genomics showed that several verified SNPs affected evolutionarily constrained parts of the mitochondrial genome. Two approaches were used, blasting salmon EST’s to NCBI (May 2013) and retrieving multiz alignments of vertebrates from the UCSC genome browser (in September 2013). This yielded several hundred sequences from related fish and other vertebrates. The list was reduced to 20 sequences, aligned with ClustalW and manually adjusted in Genedoc. The species and genome versions used are; Human (*Homo sapiens*, hg19), Lamprey (*Petromyzon marinus*, petMar1), Fugu (*Takifugu rubripes*, fr2), Medaka (*Oryzias latipes*, oryLat2), Stickleback (*Gasterosteus aculeatus*, gasAcu1), Tetraodon (*Tetraodon nigroviridis*, tetNig2), Zebrafish (*Danio rerio*, danRer6). We also downloaded from NCBI the sequence of whole or partial mtDNA from several fish species; Brown trout (*Salmo trutta*, JQ390057 and AF148843), Broad whitefish (*Coregonus nasus*, JQ390058), Legless searsid (*Platytroctes apus*, AP004107), Pacific menhaden (*Ethmidium maculatum*, AP011602), Icefish (*Salanx ariakensis*, AP006231 and HM151535), Chain pickerel (*Esox niger*, AP013046) and Western Pacific roughy (*Hoplostethus japonicus*, AP002938). The three mitochondrial variants (numbered by the *S. alpinus* mtDNA - NC_000861) are; ml829G>A (CCACGTTGTGAAACCAAC[G/A]TCCGAAGGTGGATTTAGCAGT), m3211T>C (CGTGCAGAAGCGGGCATAAG[T/C]ACATAAGACGAGAAGACCCT) and m3411C>T (CTCTAAGCACCAGAATTT[C/T]TGACCAAAAATGATCCGGC). The images were produced in GIMP version 2.8.6. for Windows [81].

## Results

### RNA sequencing characteristics

Our long term aim is to get a handle on molecular and developmental systems that relate to the frequent ecological and phenotypic divergence in Arctic charr, including craniofacial features that emerge some weeks before and after hatching of embryos. To this end we analysed transcriptomes of pooled embryonic samples at four time-points (141, 163, 200 and 433 *τs*) derived from pure crosses of two Arctic charr morphs with contrasting phenotypes; the small benthic morph from Lake Thingvallavatn (SB) and aquaculture charr (AC). Each sample yielded good quality data, with sequencing depth from 49 to 58 (average 55) million reads. To quantify the expression levels, the reads were aligned to a salmon EST-assembly [69]. Around 20% of the reads mapped uniquely to the EST data (S2 Table). A further 30% mapped to two or more contigs, probably representing paralogous genes, recent duplications or repeat like elements within transcribed regions. A substantial fraction of the RNA-sequencing reads did not map to the contigs from *S. salar* (S2 Table). Analyses of those reads require an Arctic charr genome sequence or transcriptome assembly from longer and paired end reads.

### Differential expression during Arctic charr development

The scope of the sampled developmental timepoints and morphs only allows contrasts of expression between morphs, or expression differences between time points. For the expression analysis ESTs were collapsed into 16851 genes or paralog groups (“gene” has here this broader meaning, see Materials and Methods). We only considered genes (total of 10496) with 800 or more reads mapped and tested for differential expression using the edgeR-package [71].

We detected considerable changes in the transcriptome during Arctic charr development (Figure 2A). The expression of 1603 and 2459 genes differed significantly between developmental timepoints at the 1% and 5% levels of false discovery rate (FDR), respectively (S1 File). The difference was most pronounced between pre-hatching (timepoints: 141, 163, 200 *τs*) and post hatching embryos (timepoint 433 *τs*), as more than 70% of the genes with FDR below 1% had higher expression in the latter (Figure 2A). According to Gene Ontology analyses, six separate GO categories are significant (below 10%FDR). The most drastic changes were seen in processes related to glycolysis (GO:0006096, FDR = 0.0009), were the expression of 19 out of 25 genes changed during this developmental period. The other five classes that were differentially expressed during charr development are, ion transport (GO:0006811, FDR = 0.027), blood coagulation (GO:0007596, FDR = 0.03), DNA repair (GO:0006281, FDR = 0.08) and two immune related categories (GO:0019882, FDR = 0.08, GO:0006955, FDR = 0.09). Those results probably reflect developmental changes and/or differences in the environment of embryos before and after hatching.

**Figure 2. Heatmap of differentially expressed genes in the Arctic charr developmental transcriptome.** Two morphs (SB and AC) are represented, at four timepoints. (A) The 1603 genes with expression difference among time points, here clustered into four groups. (B) The 71 genes differentially expressed between morphs, are clustered into 4 groups for each of the two morphs. High expression is indicated by blue and low expression by beige.

### Differential expression between Arctic charr morphs

We were especially interested in genes showing expression differences between the two Arctic charr morphs as such differences may relate to pathways involved in phenotypic divergence. In the data 296 genes were differentially expressed (FDR < 5%) between the morphs (141 higher in SB and 152 higher in AC). Among genes with higher expression in SB-charr two biological GO categories were enriched: blood coagulation (GO:0007596, p = 0.001) and proteolysis (GO:0006508, p = 0.002). Recall, expression of blood coagulation factors also differed between developmental stages (see above). In AC-charr, genes in three categories: respiratory electron transport chain (GO:0022904, p = 0.0006), ATP synthesis coupled electron transport (GO:0042773, p = 0.002) and neurotransmitter transport (GO:0006836, p = 0.009) have higher expression. The first two GO categories both relate to energy generation in mitochondria and may imply higher expression of genes with mitochondrial functions in AC-charr.

Using more stringent FDR (1%), 31 genes had higher expression in SB and 40 genes higher in AC (Figure 2B, Tables 1 and 2). These genes have diverse functional annotations. The genes with higher expression in each morph were clustered into 4 groups, which aggregated genes of similar function. For instance SB cluster 3 has three immune related genes: *Complement factor D* (9), *H-2 class I histocompatibility antigen L-D alpha chain* (2) and *Sushi domain-containing protein 2* (4) and one gene with unknown function (Table 1). Note, however, that immune genes were not significantly enriched in the GO comparison of morphs. Testing for heterochrony in expression was not feasible, as only one sample per morph and developmental time was sequenced.

**Table 1.** Differentially expressed genes, with higher expression in the SB morph from Lake Thingvallavatn.

| NR | Unigene.Description | NR.contigs | logFC | logCPM | FDR | Cluster |
| --- | --- | --- | --- | --- | --- | --- |
| 3766 | Histone H3 embryonic | 1 | 8.71 | 2.74 | 7.80E-035 | S-1 |
| 5103 | Natterin-like protein | 6 | 2.75 | 7.12 | 7.76E-007 | S-2 |
| 356 | A7J6M9 Putative uncharacterized protein n175R | 1 | 2.33 | 4.66 | 3.30E-006 | S-1 |
| 6697 | Q1KY05 Main olfactory receptor-like protein | 5 | 3.12 | 6.92 | 9.96E-005 | S-1 |
| 8151 | Sushi domain-containing protein 2 | 4 | 2.20 | 5.55 | 0.0001 | S-3 |
| 1682 | Carcinoembryonic antigen-related cell adhesion molecule 1 | 3 | 2.55 | 3.83 | 0.0002 | S-1 |
| 6228 | Protein FAM98A | 2 | 1.96 | 4.76 | 0.0003 | S-1 |
| 7531 | STAM-binding protein-like | 2 | 2.07 | 2.62 | 0.0005 | S-1 |
| 6712 | Q1M160 Myc-regulated DEAD box protein | 1 | 1.67 | 3.23 | 0.0009 | S-1 |
| 2300 | Cytosolic sulfotransferase 3 | 3 | 1.73 | 2.13 | 0.0009 | S-1 |
| 2063 | Complement factor D | 7 | 1.79 | 6.42 | 0.0016 | S-3 |
| 3326 | Galectin-3-binding protein A | 4 | 1.79 | 3.85 | 0.0017 | S-4 |
| 3169 | Flocculation protein FLO11 | 2 | 1.86 | 4.05 | 0.0017 | S-1 |
| 1203 | B5XDY0 H-2 class I histocompatibility antigen L-D alpha chain | 2 | 1.70 | 2.12 | 0.0028 | S-3 |
| 9183 | UPI000065D844 UPI000065D844 related cluster | 2 | 1.97 | 5.55 | 0.0028 | S-1 |
| 2909 | Epidermis-type lipoxigenase 3 | 4 | 1.68 | 4.84 | 0.0029 | S-1 |
| 4884 | Myeloperoxidase | 4 | 2.20 | 6.78 | 0.0029 | S-1 |
| 10003 | Uridine phosphorylase 1 | 4 | 1.51 | 3.00 | 0.0047 | S-1 |
| 2513 | Desmoglein-1-alpha | 1 | 1.59 | 2.80 | 0.0054 | S-2 |
| 377 | A7SJA8 Predicted protein (Fragment) | 1 | 1.73 | 2.50 | 0.0055 | S-3 |
| 9204 | UPI00006A2900 UPI00006A2900 related cluster | 2 | 6.38 | 3.26 | 0.0064 | S-1 |
| 9642 | UPI00017B1B0F UPI00017B1B0F related cluster | 1 | 2.00 | 1.92 | 0.0064 | S-2 |
| 1965 | Coiled-coil domain-containing protein 136 | 2 | 2.15 | 2.32 | 0.0064 | S-2 |
| 9260 | UPI0000F1D4BA PREDICTED: hypothetical protein | 1 | 1.80 | 2.41 | 0.0065 | S-2 |
| 738 | Adseverin | 8 | 1.58 | 5.51 | 0.0073 | S-1 |
| 9678 | UPI00017B4479 UPI00017B4479 related cluster | 1 | 2.18 | 1.97 | 0.0074 | S-4 |
| 8339 | Testin | 4 | 1.50 | 4.93 | 0.0080 | S-2 |
| 6840 | Q4SNH3 Chromosome 8 SCAF14543 whole genome shotgun sequence | 1 | 1.42 | 4.00 | 0.0080 | S-1 |
| 1668 | Carbohydrate sulfotransferase 6 | 1 | 2.09 | 2.08 | 0.0090 | S-4 |
| 8341 | Testisin | 2 | 2.01 | 2.76 | 0.0090 | S-4 |
| 6373 | Protein asteroid homolog 1 | 6 | 1.29 | 4.24 | 0.0090 | S-4 |

logFC – log Fold Change
logCPM – log Counts Per Million
FDR – False Discovery Rate
The cluster numbering corresponds to Figure 1.

**Table 2.** Differentially expressed genes, with higher expression in the AC morph.

| NR | Unigene.Description | NR.contigs | logFC | logCPM | FDR | Cluster |
| --- | --- | --- | --- | --- | --- | --- |
| 3465 | Glutathione S-transferase P 1 | 1 | -8.35 | 2.45 | 1.12E-019 | A-2 |
| 2475 | Dehydrogenase/reductase SDR family member 7 | 2 | -4.88 | 2.15 | 9.67E-014 | A-3 |
| 6945 | Q6NWE8 Sb:cb283 protein (Fragment) | 3 | -6.08 | 3.02 | 2.15E-013 | A-2 |
| 399 | A8DW32 Predicted protein | 1 | -5.32 | 6.38 | 4.27E-010 | A-1 |
| 9682 | UPI00017B4B48 UPI00017B4B48 related cluster | 2 | -3.70 | 2.81 | 2.61E-008 | A-2 |
| 9817 | Uncharacterized protein ART2 | 5 | -12.63 | 6.89 | 8.23E-008 | A-2 |
| 6724 | Q2L0Z2 Putative ATP-dependent RNA helicase | 1 | -3.41 | 1.89 | 1.88E-007 | A-2 |
| 1197 | B5XD10 Vacuolar proton pump subunit G 1 | 1 | -4.30 | 2.10 | 1.84E-006 | A-2 |
| 5325 | Nucleoside diphosphate kinase B | 1 | -9.85 | 7.63 | 2.51E-006 | A-1 |
| 9205 | UPI0000D5B923 PREDICTED: myelin basic protein isoform 1 | 3 | -2.49 | 3.45 | 9.18E-006 | A-3 |
| 6377 | Protein broad-minded | 1 | -2.11 | 2.74 | 4.75E-005 | A-1 |
| 5711 | Pistil-specific extensin-like protein | 1 | -2.16 | 2.60 | 0.0002 | A-3 |
| 3203 | Formin-like protein 20 | 7 | -1.98 | 1.95 | 0.0002 | A-3 |
| 9315 | UPI0000F2EC69 PREDICTED: hypothetical protein | 2 | -5.60 | 4.57 | 0.0005 | A-2 |
| 363 | A7RFV0 Predicted protein (Fragment) | 2 | -1.74 | 4.96 | 0.0010 | A-3 |
| 6937 | Q6AZT1 MGC81677 protein | 3 | -2.06 | 3.81 | 0.0014 | A-2 |
| 3756 | Histone H1 | 3 | -2.26 | 4.54 | 0.0017 | A-2 |
| 1133 | B5DGN9 Creatine kinase-1 | 7 | -4.72 | 5.50 | 0.0017 | A-3 |
| 309 | A1IMH7 CD80-like protein | 12 | -1.94 | 4.29 | 0.0017 | A-2 |
| 7651 | Serine protease ami | 2 | -1.54 | 5.90 | 0.0017 | A-3 |
| 9935 | Uncharacterized protein C7orf63 homolog | 1 | -1.87 | 1.91 | 0.0025 | A-2 |
| 5219 | Nostrin | 2 | -2.55 | 3.38 | 0.0029 | A-2 |
| 1855 | Chondroitin sulfate N-acetylgalactosaminyltransferase 2 | 5 | -2.56 | 6.14 | 0.0034 | A-1 |
| 10203 | Xylose isomerase | 6 | -1.55 | 2.43 | 0.0035 | A-3 |
| 2249 | Cytochrome c oxidase subunit 3 | 11 | -1.78 | 11.15 | 0.0035 | A-3 |
| 180 | 40S ribosomal protein S3-B | 2 | -5.31 | 8.67 | 0.0050 | A-1 |
| 1227 | B6NBL3 Putative uncharacterized protein | 3 | -1.59 | 2.95 | 0.0050 | A-2 |
| 5055 | NADH-ubiquinone oxidoreductase chain 6 | 2 | -1.48 | 2.65 | 0.0061 | A-3 |
| 4634 | Metallothionein A | 1 | -3.33 | 5.44 | 0.0064 | A-2 |
| 342 | A5C0J4 Putative uncharacterized protein | 2 | -2.58 | 2.47 | 0.0064 | A-2 |
| 9698 | UPI00019258B4 PREDICTED: similar to epithelial cell transforming sequence 2 oncogene protein partial | 1 | -2.06 | 2.94 | 0.0064 | A-2 |
| 5878 | Pro-opiomelanocortin B | 1 | -2.04 | 5.60 | 0.0065 | A-2 |
| 2248 | Cytochrome c oxidase subunit 2 | 9 | -2.21 | 9.83 | 0.0074 | A-2 |
| 1246 | B8JI87 Novel protein similar to vertebrate collagen type VI alpha 3 (COL6A3) (Fragment) | 1 | -1.69 | 3.16 | 0.0080 | A-3 |
| 7994 | Sperm-associated antigen 5 | 1 | -2.07 | 3.71 | 0.0080 | A-2 |
| 9515 | UPI000175F90F PREDICTED: similar to pleckstrin homology domain containing family A member 7 | 1 | -2.00 | 1.87 | 0.0090 | A-2 |
| 1124 | B5DDZ4 Acta1 protein | 1 | -1.52 | 2.62 | 0.0090 | A-2 |
| 1127 | B5DG94 2-peptidylprolyl isomerase A | 2 | -2.56 | 5.67 | 0.0090 | A-1 |
| 9175 | UPI000054A3C0 PREDICTED: apolipoprotein B | 3 | -1.32 | 4.09 | 0.0090 | A-4 |
| 9671 | UPI00017B3C62 UPI00017B3C62 related cluster | 1 | -1.51 | 1.92 | 0.0096 | A-1 |

logFC – log Fold Change
logCPM – log Counts Per Million
FDR – False Discovery Rate
The cluster numbering corresponds to Figure 1.

The results suggest mitochondrial function and blood coagulation genes are differentially expressed between the morphs, but due to few samples used in the RNA-sequencing, qPCR verification was needed.

### Validation of gene expression differences in whole embryos and paralog specific expression of natterin genes

The transcriptome data highlights genes likely to differ in expression between embryos of SB- and AC-charr. Nine were chosen for validation by qPCR in whole embryos. Of those, five genes were confirmed to be differentially expressed between AC and SB at 161 or 200 *τs* (Figure 3, S3 Table). Three of these genes *Nattl, Alkaline phosphatase (Alp)* and *Lysozyme (Lyz)*, had significantly higher expression in SB. The other two, *Keratin-associated protein 4–3 (Krtap4–3)* and *Poly polymerase 6 (Parp6)* had higher expression in AC embryos (Figure 3, S3 Table). No morph and time interaction was detected for any of the genes.

**Figure 3. qPCR validation of candidates from transcriptome in whole embryos of Arctic charr.** Relative expression of 9 genes (A-I) analysed by qPCR in the small benthic (SB) charr from Lake Thingvallavatn and aquaculture (AC) charr at two different developmental timepoints (161 and 200 *τs).* 5 genes were differentially expressed between the two morphs *(Alp, Krtap4-3, Lyz, Nattl, Parp6)*, while 4 further genes did not show significant expression differences between morphs *(Cgat2, Cox6B1, Ndub6, Ubl5)*, see S3 Table. Error bars represent standard deviation calculated from two biological replicates.

As some of the genes are represented by different contigs or even paralogs, we set out to disentangle the expression of one paralog group in detail. The qPCR primers used above matched conserved gene regions and thus are likely to estimate the combined expression of several paralogs. We chose to measure the expression of three different *natterin* paralogs (*nattl 1, 2 and 3*), in part because this understudied gene was first characterized as a toxin produced by a tropical fish [82, 83]. We studied *nattl* expression in several developmental stages in AC-, SB- and PL-charr as well as in selected tissues of adult AC-charr. The expression level of the three paralogs differed between morphs and timepoints (Figure 4 and S4 Table). Overall *nattl2* had the highest expression in all morphs. The *nattl1* had higher expression in embryos of PL-charr than in AC- and SB-charr, while *nattl2* and *nattl3* were more expressed in SB-embryos.

**Figure 4. Relative expression of *Nattl* and its three paralogs during charr development in different morphs.** The expression is graphed for different morphs (SB, AC and PL) at four developmental timepoints (161, 200, 256 & 315 *τs*, relative to AC-charr at timepoint 161. A) General *Nattl* expression along charr development. B-D) Expression of *Nattl paralogs* 1-3. ANOVA showing the variation among morphs is summarized in S4 Table.

In order to evaluate the hypothesis that *nattl* genes have immune-related functions we studied expression in adult tissues (in AC-charr). The *nattl* expression was highest in the gills, followed by expression in kidney, skin and spleen. Low expression levels were detected in liver, intestine and heart (S1 Fig and S4 Table). The three *nattl* paralogs followed different patterns, whilst each of them showed significant expression differences among tissues. *Nattl1* was mainly expressed in spleen and kidney, while *nattl2* showed a significantly higher expression in skin, liver and in gills. Similarly, the relative expression of *nattl3* was highest in the gills and skin. This indicates that the three *nattl* paralogs are expressed in a tissue specific manner, and also differently during the development of the three charr morphs studied here.

In sum, we confirm the differential expression of five of nine genes between the morphs. Furthermore, *nattl* paralogs have tissue specific expression pattern in adults, that may reflect immunological functions.

### Expression differences in the developing heads of benthic and limnetic charr morphs

To study the craniofacial divergence between sympatric Arctic charr morphs we used qPCR to study 8 genes with expression difference in the RNA-seq data (all higher in SB). We focused on genes with known craniofacial expression in zebrafish development [84] and compared two benthic (SB, LB) and two limnetic charr (AC, PL). We analyzed heads at three time-points (178, 200 and 218 *τs*) as this period overlaps with early stages of craniofacial skeletal formation in Arctic charr [85, 86]. The qPCR confirmed the higher expression of seven out of these eight genes in the head of benthic charr compared to limnetic charr (Figure 5 and S2 Fig). These seven genes are *Claudin 4 (Cldn4), adseverin (Scin), Junction plakoglobin (Jup), Lipolysis stimulated lipoprotein receptor (Lsr), Major vault protein (Mvp), Transforming growth factor beta receptor II (Tgfbr2)* and *Vitamin D receptor a (Vdra).* The eighth gene, *Retinoic acid receptor gamma-A (Rarg)* gave a small but significant response in the head, but the effects were reversed, i.e. the expression was higher in AC. The expression difference of the seven genes was, in almost all cases, consistent over the three time-points studied (See S2 Fig). These results show that RNA-sequencing of Aquaculture charr with limnetic craniofacial morphology and small benthic charr can be used to reveal differential expression of genes that associate with limnetic and benthic divergence in craniofacial elements in sympatric charr morphs.

**Figure 5. Expression differences of craniofacial candidate genes in developing head of Arctic charr morphs.** Relative expression ratios, calculated from the qPCR data, were subjected to an ANOVA to test the expression differences amongst four charr groups and three close time points (*τs*). The underlined gene names reflect significant difference between SB and AC-charr. A post hoc Tukey’s test (HSD) was performed to determine the effects of morphs, time and morph-time interaction (M X T). White boxes represent low expression, while black boxes represent high expression. The shading represents significant different expression between the samples (*α* = 0.05, NS = not significant).

### Analyses of polymorphism in Arctic charr transcriptome

The RNA-seq data also revealed segregating variations with large frequency differences between charr morphs. To uncover candidate SNPs we mapped the reads to all of the *S. salar* EST-contigs. Filtering on coverage yielded 165,790 candidate SNPs (Table 3); of those 66.569 came from reads that mapped uniquely (UNI) and 57.009 candidate SNPs from reads that mapped to more than one contig (REP); with limited overlap between lists. The salmonid ancestor underwent whole genome duplication, generating long blocks of synteny and paralogous genes [36]. Assuming that the expression of paralogous genes is stable, then differences among paralogs appear as SNPs at similar frequency in all samples. By requiring variant frequency differences (p < 0.05, uncorrected) between samples we reduced the list of candidates by two thirds, yielding over 20.000 candidate SNPs. Note, that as cDNA from charr families was sequenced (not a population sample), estimates of SNP frequencies are imprecise. To err on the side of caution, we chose SNP candidates with 50% or higher frequency difference between morphs for further study. The candidate SNPs were also summarized by frequency of the derived allele, in reference to the *S. salar* sequence. This gave 672 and 872 SNPs at higher frequency, in AC-charr and SB-charr, respectively. The uniquely and multiply mapped reads, revealed approximately similar numbers of candidate SNPs. Gene ontology analysis showed that for derived SNPs in SB, there was an excess of variants in genes related to translation, both as a broad category and specific subgroups (S5 Table). There was also enrichment of SNPs in genes related to DNA-mediated transposition, DNA integration, DNA replication and oxidation-reduction process. No GO categories were enriched for high frequency derived SNPs in AC. Furthermore, functional effects of the candidate SNPs (UTR, synonymous and non-synonymous) were predicted. The distribution among those categories did not differ between variants detected by uniquely or repeatedly mapped reads, 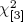 = 2.59, *p* = 0.46 (S6 Table).

**Table 3.** Candidate SNPs in the Arctic charr transcriptome, filtered by coverage, difference between sample and morphs and frequency difference between morphs. For Delta > 0.95 we show the number of SNP-candidates before the redundant ones were removed.

| SNP-candidates | Morph | Uni | Rep | Total |
| --- | --- | --- | --- | --- |
| Total |  | 96231 | 74341 | 165790 |
| Filter coverage |  | 66569 | 57009 | 113776 |
| Diff. Bwn. samples |  | 21417 | 22252 | 42869 |
| Diff. Bwn. morphs |  | 11385 | 12953 | 23974 |
| Delta > 0.5 | AC | 396 | 285 | 672 |
| Delta > 0.5 | SB | 526 | 353 | 872 |
| Delta > 0.75 | AC | 95 | 68 | 159 |
| Delta > 0.75 | SB | 155 | 95 | 248 |
| Delta > 0.95 | AC | 17 | 13 | 30 |
| Delta > 0.95 | SB | 29 | 4 | 33 |

SNP-candidates are found by mapping to *S. salar* ESTs. From UNIquely or REPeatedly mapped RNA-reads. Delta: Differences in allele frequency between morphs, categorized by which morph had the higher derived allele frequency.

A total of 60 candidate SNPs are nearly fixed in one morph, with frequency difference between morphs above 95% (after manual inspection of contigs and SNP position three candidates were removed since they represented the same SNP). Of these “fixed” SNPs 46 came from uniquely mapped reads and 14 from reads that mapped more than twice (Table 4 and 5). For the SNPs from uniquely mapped reads, 17 are fixed in AC-charr and 29 in SB-charr. The few genes with two or more polymorphic sites were; *Keratin type II cytoskeletal 3 (KRT3), Cysteine sulfinic acid decarboxylase (CSAD)* and *DNA-directed RNA polymerase I subunit RPA12 (RPA12)* with 5, 5 and 2 SNPs respectively. *KRT3* and *CSAD* had significant differentiation in both SB and AC. Similarly, 14 SNPs with large differentiation between morphs were predicted from reads that mapped on two or more contigs (Table 5). Of these, we found two variants in the mitochondrial *60S ribosomal protein L36 (RPL36)* and variants in 4 other mitochondrial genes (*28S ribosomal protein S18a mitochondrial (MRPS18A), Apoptosis-inducing factor 1 mitochondrial (AIFM1), Isocitrate dehydrogenase [NADP] mitochondrial (acIDH1)* and *Protein S100-A1 (S100A1))*, all at higher frequency in AC-charr. PCR and Sanger sequencing was used to confirm SNPs in *DNA2-like helicase (DNA2)*, a gene with nuclear and mitochondrial function, and two other genes *Uroporphyrinogen decarboxylase (UROD)*, and *Mid1-interacting protein 1-like (MID1IP1)* (S7 Table). The candidate variant *Eukaryotic translation initiation factor 4 gamma 2 (EIF4G2)* was not substantiated by the PCR/sequencing.

**Table 4.**
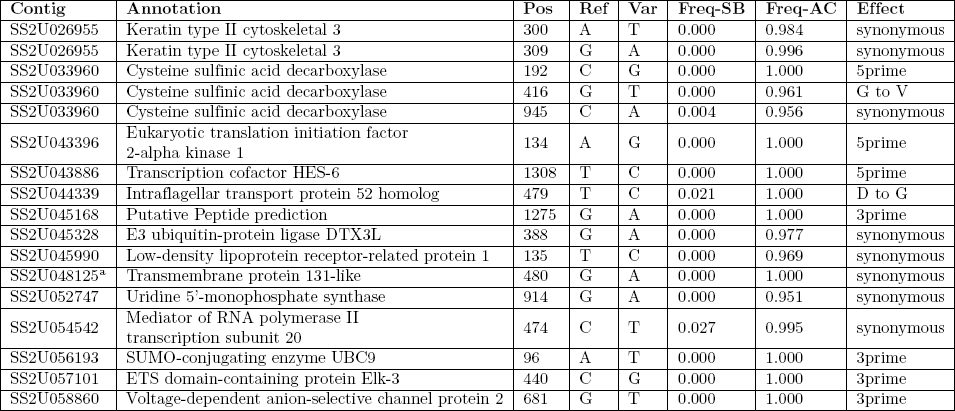
SNP candidates from uniquely mapped reads. **(a)** Higher frequency in AC morph

| Contig | Annotation | Pos | Ref | Var | Freq-SB | Freq-AC | Effect |
| --- | --- | --- | --- | --- | --- | --- | --- |
| SS2U026955 | Keratin type II cytoskeletal 3 | 300 | A | T | 0.000 | 0.984 | synonymous |
| SS2U026955 | Keratin type II cytoskeletal 3 | 309 | G | A | 0.000 | 0.996 | synonymous |
| SS2U033960 | Cysteine sulfinic acid decarboxylase | 192 | C | G | 0.000 | 1.000 | 5prime |
| SS2U033960 | Cysteine sulfinic acid decarboxylase | 416 | G | T | 0.000 | 0.961 | G to V |
| SS2U033960 | Cysteine sulfinic acid decarboxylase | 945 | C | A | 0.004 | 0.956 | synonymous |
| SS2U043396 | Eukaryotic translation initiation factor 2-alpha kinase 1 | 134 | A | G | 0.000 | 1.000 | 5prime |
| SS2U043886 | Transcription cofactor HES-6 | 1308 | T | C | 0.000 | 1.000 | 5prime |
| SS2U044339 | Intraflagellar transport protein 52 homolog | 479 | T | C | 0.021 | 1.000 | D to G |
| SS2U045168 | Putative Peptide prediction | 1275 | G | A | 0.000 | 1.000 | 3prime |
| SS2U045328 | E3 ubiquitin-protein ligase DTX3L | 388 | G | A | 0.000 | 0.977 | synonymous |
| SS2U045990 | Low-density lipoprotein receptor-related protein 1 | 135 | T | C | 0.000 | 0.969 | synonymous |
| SS2U048125<sup>a</sup> | Transmembrane protein 131-like | 480 | G | A | 0.000 | 1.000 | synonymous |
| SS2U052747 | Uridine 5'-monophosphate synthase | 914 | G | A | 0.000 | 0.951 | synonymous |
| SS2U054542 | Mediator of RNA polymerase II transcription subunit 20 | 474 | C | T | 0.027 | 0.995 | synonymous |
| SS2U056193 | SUMO-conjugating enzyme UBC9 | 96 | A | T | 0.000 | 1.000 | 3prime |
| SS2U057101 | ETS domain-containing protein Elk-3 | 440 | C | G | 0.000 | 1.000 | 3prime |
| SS2U058860 | Voltage-dependent anion-selective channel protein 2 | 681 | G | T | 0.000 | 1.000 | 3prime |

**Table 4.** SNP candidates from uniquely mapped reads. **(b)** Higher frequency in SB morph

| Contig | Annotation | Pos | Ref | Var | Freq-SB | Freq-AC | Effect |
| --- | --- | --- | --- | --- | --- | --- | --- |
| SS2U000399 | Insulin-like growth factor-binding protein 7 | 598 | C | A | 1.000 | 0.000 | 3prime |
| SS2U004484 | Titin | 387 | G | A | 0.990 | 0.010 | synonymous |
| SS2U026826 | L-asparaginase | 363 | C | T | 1.000 | 0.000 | H to Y |
| SS2U026955 | Keratin type II cytoskeletal 3 | 116 | C | A | 0.996 | 0.031 | T to N |
| SS2U026955 | Keratin type II cytoskeletal 3 | 264 | C | T | 0.970 | 0.008 | synonymous |
| SS2U026955 | Keratin type II cytoskeletal 3 | 317 | C | T | 1.000 | 0.002 | T to M |
| SS2U033960 | Cysteine sulfinic acid decarboxylase | 363 | C | T | 1.000 | 0.025 | 5prime |
| SS2U033960 | Cysteine sulfinic acid decarboxylase | 387 | C | T | 1.000 | 0.030 | synonymous |
| SS2U033960 | Cysteine sulfinic acid decarboxylase | 657 | T | C | 0.990 | 0.031 | synonymous |
| SS2U034322 | Cyclin-C | 1094 | A | G | 1.000 | 0.000 | 3prime |
| SS2U034431 | Dolichyl-diphosphooligosaccharide-protein glycosyltransferase subunit 2 | 436 | G | A | 0.992 | 0.000 | G to S |
| SS2U036025 | Nuclear receptor coactivator 4 | 36 | G | A | 1.000 | 0.043 | 5prime |
| SS2U040590 | Glutamyl-tRNA(Gln) amidotransferase subunit A homolog | 478 | G | A | 0.972 | 0.000 | synonymous |
| SS2U045606 | Superkiller viralicidic activity 2-like 2 | 500 | C | T | 1.000 | 0.000 | synonymous |
| SS2U047816 | Squalene synthase | 1139 | G | A | 1.000 | 0.029 | synonymous |
| SS2U048063 | Lysine-specific demethylase NO66 | 669 | C | T | 1.000 | 0.000 | synonymous |
| SS2U050394 | UPF0542 protein C5orf43 homolog | 596 | G | A | 1.000 | 0.000 | synonymous |
| SS2U050880<sup>a</sup> | Transmembrane protein 131-like | 901 | C | T | 1.000 | 0.000 | A to V |
| SS2U052076 | Eukaryotic translation initiation factor 3 subunit A | 824 | C | T | 1.000 | 0.031 | synonymous |
| SS2U053417 | RNA polymerase-associated protein LEO1 | 454 | G | A | 1.000 | 0.049 | synonymous |
| SS2U054333 | Scaffold attachment factor B2 | 382 | G | A | 0.999 | 0.000 | V to M |
| SS2U054705 | Cell division protein kinase 4 | 122 | A | G | 0.971 | 0.000 | 3prime |
| SS2U054965 | DNA-directed RNA polymerase I subunit RPA12 | 106 | G | A | 1.000 | 0.000 | 5prime |
| SS2U054965 | DNA-directed RNA polymerase I subunit RPA12 | 411 | T | G | 1.000 | 0.000 | synonymous |
| SS2U055120 | Chromatin modification-related protein MEAF6 | 350 | A | C | 1.000 | 0.000 | H to P |
| SS2U055153 | Complexin-1 | 1191 | C | A | 1.000 | 0.031 | 3prime |
| SS2U057635 | Mitogen-activated protein kinase 14B | 1370 | A | T | 1.000 | 0.026 | 3prime |
| SS2U058169 | Transmembrane protein 50A | 1214 | C | G | 0.973 | 0.000 | 3prime |
| SS2U058802 | Signal recognition particle 54 kDa protein | 607 | T | A | 0.969 | 0.000 | C to S |

<sup>a</sup> Those genes are distinct paralogs

**Table 5.** SNP candidates with significant difference frequency between AC and SB morphs, from reads that mapped to two or more contigs.

| Contig | Annotation | Pos | Ref | Var | Freq-SB | Freq-AC | Effect |
| --- | --- | --- | --- | --- | --- | --- | --- |
| SS2U004839 | Actin alpha sarcomeric/cardiac | 550 | A | C | 0.015 | 0.999 | 3prime |
| SS2U021298 | 28S ribosomal protein S18a mitochondrial | 462 | A | C | 0.000 | 1.000 | synonymous |
| SS2U041264 | Apoptosis-inducing factor 1 mitochondrial | 341 | C | T | 0.000 | 0.952 | synonymous |
| SS2U054211<sup>a</sup> | Cytoplasmic dynein 1 intermediate chain 2 | 136 | T | C | 0.018 | 0.974 | synonymous |
| SS2U054362<sup>a</sup> | Q08CA8 Dynein cytoplasmic 1 intermediate chain 2 | 945 | A | G | 0.000 | 1.000 | synonymous |
| SS2U055923 | Bystin | 1623 | A | C | 0.000 | 0.983 | 3prime |
| SS2U058758 | Protein S100-A1 | 253 | C | T | 0.000 | 0.984 | synonymous |
| SS2U059000 | Isocitrate dehydrogenase [NADP] mitochondrial | 1654 | T | C | 0.000 | 0.975 | 3prime |
| SS2U059146 | 60S ribosomal protein L36 | 263 | T | G | 0.009 | 1.000 | synonymous |
| SS2U059146 | 60S ribosomal protein L36 | 470 | A | C | 0.009 | 1.000 | synonymous |
| SS2U036667 | Heterogeneous nuclear ribonucleoprotein K | 813 | C | T | 1.000 | 0.022 | 5prime |
| SS2U042873 | RNA polymerase-associated protein LEO1 | 460 | G | A | 1.000 | 0.000 | synonymous |
| SS2U058455 | Adenylosuccinate lyase | 1616 | C | T | 1.000 | 0.000 | 3prime |
| SS2U058906 | Mid1-interacting protein 1-like | 350 | G | T | 0.985 | 0.000 | E to D |

<sup>a</sup> Those genes are distinct paralogs

In sum, the data suggest substantial genetic separation between the two charr morphs studied here, the small benthic from Lake Thingvallavatn and the aquaculture charr from Holar. While individual SNPs clearly need to be verified e.g. by Sanger sequencing or SNP assays on more individuals, the results suggests genetic differentiation between these morphs in genes of various molecular systems. Considering the enrichment of differentially expressed genes related to mitochondrial energy metabolism (above), and high frequency candidate SNPs in several genes with mitochondrial function in AC-charr we decided to study the mitochondrial transcriptome further.

### Polymorphism and expression of Arctic charr mtDNA

The charr studied here reflect metabolic extremes, the aquaculture charr was bred for growth while the small benthic morph is thought to have experienced natural selection for slow metabolism and retarded growth [49, 87]. Although mRNA preparation protocols were used for generating cDNA for the RNA-sequencing, a substantial number of reads came from non-polyadenylated sequences. By mapping the reads to mtDNA sequence of Arctic charr we could estimate expression of mitochondrial genes and infer polymorphism both in genes and intergenic regions. There was a clear difference in sequencing coverage, with more than twice as many reads mapped from the AC-compared to SB-charr (mean fold difference 2.27, Wilcoxon test, p < 0.0004). Note, as only two types of fish are compared, it is impossible to determine the polarity of expression divergence.

Using an appropriate outgroup it is possible to determine ancestral and derived states for DNA polymorphism data. The mapped RNA-reads were used to identify polymorphism and divergence in the entire mitochondrial chromosome. The polymorphisms were found by mapping to mtDNA from a Canadian *S. alpinus* [61], but ancestral vs. derived status inferred by comparison to *S. salar* mtDNA. Bioinformatics revealed 82 candidate sites, including 35 that represent divergence between Icelandic and Canadian charr. A total of 20 candidate SNPs had high (more than 50%) frequency difference between SB- and AC-charr (Figure 6). There was no bias in the distribution of derived SNPs, 11 on the AC branch and 9 in SB. Note, the frequency distribution is most irregular as we sequenced embryos of related individuals (see Materials and Methods), not a population sample. The divergence between Iceland and Canada is particularly little in the 12s and 16s ribosomal RNA genes. Curiously in those genes were two SNPs differing strongly in frequency between morphs (Figure 6). To confirm and better estimate the frequency of variants in the ribosomal genes, we PCR amplified and sequenced two ~550 bp regions in the rRNA genes, comparing three morphs (PL, LB and SB) from Lake Thingvallavatn (Figure 7A, C and E). The 12s polymorphism (ml829G>A) differed significantly between the morphs (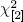 = 8.6, *p* = 0.014), and was at highest frequency in the SB (0% in PL, 12.5% in LB and 75% in SB). Similarly m3411C>T in the 16s was enriched in SB (62.5%) but found at lower frequency in PL (0%) and LB (12.5%) (it differed significantly between morphs, 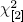 = 9.3333, *p =* 0.009). The Sanger sequencing also revealed three other polymorphisms in the amplified region, not seen in the transcriptome. Among those m3211T>C in the 16s gene was at 75% frequency in LB, but not found in the other morphs (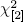 = 19.76, *p* < 0.0001).

In order to gauge the potential functionality of those variants we aligned the rRNA genes from nearly hundred fishes and several vertebrates. The position affected by m1829G>A and m3211T>C, in the 12s and 16s rRNAs, are not well conserved in fishes or vertebrates (Figure 7B and D). However m3411C>T, in the 16s rRNA, alters a position that is nearly invariant in 100 fish genomes (Figure 7F). The only exception is Pacific menhaden, which curiously also has T in this position. This region could not be aligned properly in other vertebrates. Thus m3411C>T alters a conserved position, but probably not very drastically as the introduced allele is tolerated in another fish species.

**Figure 6. Genetic divergence in the mtDNA between SB- and AC-charr.** The frequency differences between morphs of candidate SNPs estimated from the RNA-sequencing, graphed along the mtDNA chromosome. The SNPs indicate whether the derived allele is higher in SB (black dots) or AC (open circles). Sites of divergence between the Icelandic stocks and the Canadian reference sequence are indicated by triangles. The two black boxes represent the 12s (left) and 16s (right) rRNA genes, and gray boxes the 14 coding sequences.

In summary, the results indicate differentiation of mitochondrial function between the charr morphs of Lake Thingvallavatn and Icelandic aquaculture charr. Most curiously derived mutations in otherwise conserved ribosomal genes are found in SB-charr.

## Discussion

The objectives were to get a handle on genetic and molecular systems that associate with benthic morphology and more narrowly small benthic charr in Lake Thingvallavatn, Iceland as it is the best studied SB-charr population. To this end we performed transcriptome analysis contrasting the development of embryos from SB-charr and aquaculture charr. While our main focus is on SB-charr and its special features and adaptations, the data can potentially illustrate how AC-charr has responded to selection during breeding. We chose AC-charr as a point of reference for several reasons, i) limnetic like head morphology, ii) availability, and iii) because we wanted a strong contrast in this first survey of charr developmental diversity. The AC-charr proved a useful point of reference. As is outlined below, this transcriptome has already revealed differential expression of several developmental genes and regulators with differential expression between benthic and limnetic charr [64, 65]. Furthermore we previously found tight correlation of RNA-seq expression and qPCR estimates - in this very same transcriptome [64]. Furthermore, we have actually used the same morphs (AC and SB) and samples in a comparison of the developmental miRNA transcriptome – which reveal that expression of several miRNAs correlates with morph differences [68].

**Figure 7. Comparative genomics and population genetic differentiation in Arctic charr at 3 mtDNA locations.** Aligned are several fish genomes, with Lamprey or humans as outgroups, reflecting a 38 bp window around each of the 3 positions (asterix). A, C, E) Frequency of each of those variants in three Arctic charr populations from Lake Thingvallavatn (PL, LB and SB). A total of 8 individuals were genotyped from each morph, see methods. B) Alignment of variant m1829G>A in the 12s rRNA gene in fishes, using humans as an outgroup. D) Similar alignment of a 16s variant, m3211T>C and F) alignment of variant m3411C>T in the 16s rRNA gene.

### Developmental transcriptome of Arctic charr morphs

Here we generated an Illumina based RNA-seq transcriptome from four timepoints during early development, in two Arctic charr morphs (SB-charr and AC-charr). As no reference genome is available for Arctic charr, we mapped reads to *S. salar* EST-contigs [69] in order to estimate expression and identify candidate genetic polymorphisms. As many of the contigs are short or have overlapping annotations, we collapsed genes into paralogous genes when appropriate for the expression analysis. The main advantage is the reduction of the number of statistical tests (and hence increases statistical power). The downside is that paralog specific expression patterns are masked, as our qPCR results of the *natterin like* gene family show (Figure 3 and S1 Fig). Recent rainbow trout data shows most paralogs from the latest whole genome duplication event retain the same expression pattern [36] indicating that this scenario is probably uncommon; hence it is of considerable interest when two paralogs show distinct expression patterns [88]. In their analysis of Arctic charr gill transcriptome Norman *et al.* (2014, [25]) also used the Illumina sequencing technology to evaluate expression. Their reads were longer (2x100 bp) than in this study (36 bp) enabling them to assemble contigs. They did not consider the paralogs in their approach and merged contigs based on sequence identity. Thus the complexity of Arctic charr transcriptome still remains a mystery that advances in sequencing technology, assembly algorithms as well as genome sequencing of this species could aid in revealing.

Our data reflect differential deployment of several gene classes during Arctic charr development. Studies in salmonids and other fish have demonstrated large changes in expression during early development, including coordinated changes in many cellular and developmental systems [21, 28, 89–91]. Several blood coagulation factors genes showed significant changes during charr development, and were also more highly expressed in the SB-charr. This might reflect differences in the rate of development of blood composition, or tissue composition, in the two morphs. But further work is needed to confirm those patterns.

### Higher expression of lyzozyme and natterin-like in SB-charr

The genetic separation in two immunity genes among sympatric morphs in Lake Thingvallavatn [57] prompted us to examine the expression of *Lyz* and *nattl.* Both genes are expected to be involved in immune defenses and had higher expression in SB. The substrate of lysozyme [92] is the bacterial cell wall peptidoglycan and it acts directly on Gram-positive bacteria [93]. Lysozyme also promotes the degradation of the outer membrane and therefore indirectly acts also on Gram-negative bacteria [94]. Thus, lysozyme plays an essential role in immune defense in most eukaryotes. Another gene that caught our attention was *natterin-like.* Natterins were first discovered from the venom gland of the tropical toxic fish species *Thalassophryne nattereri* [82, 83], and are found by sequence similarity in e.g. zebrafish, Atlantic salmon and here in Arctic charr. The predicted Natterin proteins contain a mannose-binding lectin-like domain (Jacalin-domain) and a pore-forming toxin-like domain and can cause edema and pain due to kiniogenase activity [82]. Jacalin is known to stimulate functions of T- and B-cells and therefore can play an important role for immune system functions [95]. Mannose-binding lectins are pathogen recognition proteins (antibodies) and therefore are important for the acute phase response of fish [96, 97]. Our data suggest an immune related function of *nattl* genes in charr, as the highest expression was found in skin and kidney. But this needs to be studied in more detail. It is possible that higher expression of those two genes in SB-charr might reflect preparation of juveniles for bottom dwelling habitats, which may be rich in bacteria or otherwise more challenging for their immune system.

In this study we collapsed contigs into gene or paralog groups for the transcriptome analyses. The disadvantage of this approach is that differential expression in one paralog, can be masked by other related genes that do not differ between groups or have contrasting expression patterns. We looked at this by studying the expression of three paralogs of the *natterin like* genes in different morphs during Arctic charr development, and among tissues of adult AC-charr. The data show that the three *nattl* genes are expressed differentially between the morphs, thus it is not divergence in the expression of one paralog that explains the general *nattl* expression disparity in the transcriptome. Certainly, other scenarios could apply to other genes in the transcriptome, and the relative contribution of different paralogs should be evaluated systematically with deeper RNA-sequencing and longer reads.

### Expression divergence in craniofacial genes in benthic morphs

A study of the skulls of charr post-hatching embryos and juveniles from Lake Thingvallvatn, showed that some elements of the developing head ossified earlier in SB-charr than in PL-charr [98]. Our new data also demonstrate differences in craniofacial elements between AC- and SB-charr, along a limnetic vs. benthic axis [86]. Based on those differences between benthic and limnetic charr, we investigated further genes with roles in craniofacial development that were differentially expressed in the transcriptome. Guided by this transcriptome we had already found two extra-cellular matrix (ECM) remodeling genes, *Mmp2* and *Sparc* and a conserved co-expression module of genes with known roles in craniofacial morphogenesis, to have higher expression in developing heads of benthic Arctic charr morphs than in limnetic morphs [64, 65]. Bioinformatic and qPCR analyses suggest the co-expression module may potentially be affected by quantity of the transcription factor *ETS2.* These studies and the current data confirm the utility of the contrasting developmental transcriptomes for identifying candidate genes with differential expression during head development, as 7 out of 8 candidates were confirmed by qPCR. These genes had consistently higher expression in the developing head of two benthic morphs (SB and LB), and lower in fish with more limnetic phenotypes (AC and PL). This is quite striking, as three of the morphs (SB, LB and PL) studied here are of single origin and live in sympatry in Lake Thingvallavatn [57].

We focused on a few targets of Tgf-*β* and Ahr signaling pathways because of their role in craniofacial morphogenesis and transcriptional connection [99–101]. *Adseverin (Sein)* was one of the top differentially expressed genes (Table 1) and has roles in rearrangements of the actin cytoskeleton, chondrocyte differentiation and skeletal formation [102, 103]. In mice, *Scin* is demonstrated as a direct target of the Ahr pathway [104] and our data shows higher expression of *Scin* in the developing head of benthic charr embryos. Also, in the transcriptome *Lsr, Cldn4* and *Tgfbr2* had higher expression in SB-charr, and we show that higher expression of those genes associated with the benthic morphotype. *Lsr* is a molecular component of tri-cellular tight junctions [105] and has been shown to be suppressed upon Tgf-*β*1 stimulation [106] in a human cell line. Similarly, *Cldn4*, a tight junction protein with unknown role during embryonic morphogenesis, is also a common target of the Tgf-*β* and Ahr signaling pathways [107, 108]. Finally, the expression of *Tgfbr2*, encoding a receptor of Tgf-*ß* was slightly but significantly higher in the head of benthic morphs. Previous studies suggest a crucial role of *Tgfbr2* in craniofacial morphogenesis [109].

We also confirmed differential expression of other genes, including two with higher expression in SB-charr. *Mvp* is the predominant component of cytoplasmic ribonucleoprotein structures called vaults [110], which is highly conserved across eukaryotes. The vaults have been something of an enigma, but are implicated in several processes from signal transmission and immune response [111]. The *Jup* gene also showed higher expression in SB-charr. Finally, higher expression of *Vdra*, encoding the vitamin D receptor A, was found in the heads of benthic forms. The receptor regulates mineral homeostasis, osteoblast differentiation and bone metabolism [112]. We did not confirm the differential expression of *Rarg* between AC and SB, and the expression difference in the head was opposite to the transcriptome data for the whole embryos. This might be because the expression in head is masked by opposite differences in other tissue(s) in the RNA-seq data.

To summarize, we found several genes to be differentially expressed in Icelandic benthic and limnetic charr. Some of these genes belong to signaling pathways involved in bone mineralization and craniofacial morphogenesis (see above). It would be most interesting to see if expression of the same genes associates with benthic morphology from other lakes and habitats, or even in other species with similar trophic diversity.

### Energy metabolism tuning under domestication or adaptation in SB-charr?

By comparing AC and SB-charr, that represents a small benthic resource morph that has evolved repeatedly in Icelandic stream and pond habitats [56], we hoped to implicate genes and pathways involved in adaptation to these special habitats. The data point to differences between SB and AC-charr in systems related to energy metabolism, as may be expected considering their contrasting life histories. First, there is 2X higher expression of respiratory electron transport chain components in AC compared to SB-charr and 100% more mitochondrial derived reads are found in the AC-charr samples. Note that the direction of divergence is unknown, i.e. whether expression was up in AC or down in SB. Second, many derived candidate-SNPs in genes related to mitochondrial function were at high frequency on the AC branch. For instance in *S100A1*, which has been implicated in mitochondrial regulation in cardiac tissue in humans [113], but its expression is probably not exclusive to this tissue. Third, while the mitochondrial ribosomal genes generally evolve slowly, we do see derived variants at high frequency in the SB and large benthic charr in Lake Thingvallavatn. Specifically, m3411C>T in SB affects a position that is highly conserved among fish, and could affect function of the 16s rRNA. Earlier studies of mitochondrial markers in *S. alpinus* did not find large signals of divergence within Iceland [52, 54, 57], probably because they studied other genes. In summary, the results suggest divergence in mitochondrial function, due to the domestication of aquaculture charr and/or possibly reflecting adaptation of the small benthic charr in Lake Thingvallavatn.

The mitochondrion is more than a powerhouse, it integrates for instance metabolism, cell cycle and apoptosis [114]. The number of mitochondria and its functions are known to correlate with environmental attributes. For instance in Antarctic fishes under extreme cold, higher numbers of mitochondria are found in muscle and heart cells [115]. Furthermore, illustrating long-term evolutionary pressure at extreme conditions, genes with mitochondrial functions are more likely to be duplicated in such species [116]. The data presented here suggest an expression difference between morphs that could reflect differences in total number of mitochondrion, the number of mtDNA copies per mitochondrion or cell, or difference in RNA expression from the mtDNA, possibly due to evolution of mtDNA related to diet and/or temperature [117]. Further work is needed to map out the expression differences of mitochondrial related genes in more SB and anadromous charr morphs (representing more ancestral charr). The genetic signals should also be investigated by comparison of more populations and/or along ecological clines (e.g. temperature) or with respect to life history, similar to the work of Teacher *et al.* (2012, [118]).

### Conclusion

The data presented here set the stage for future investigations of the molecular and developmental systems involved in the development and divergence of the highly polymorphic and rapidly evolving Arctic charr. The results suggest genetic and expression changes in multiple systems are related to divergence among populations. The data also demonstrate differential expression between morphs in two immunological genes, possibly preparing embryos for distinct habitats, and of craniofacial developmental genes that may sculpture benthic vs. limnetic head morphology. The genetic data suggest (among other things) differentiation in the mtDNA and its functions, but larger scale and more detailed studies are needed to evaluate this and other hypotheses. Our interest is in the products of natural selection, as expression differences between groups may reflect genetic changes, e.g. in regulatory sequences or post transcriptional modifiers, but they could also reflect drift or reverberations in developmental cascades [119]. This project focused mainly on the unique small benthic charr, typically found in cold springs and small pond habitats in Iceland, particularly those with lava substratum [40, 56]. The availability of charr populations at different stages of divergence sets the stage for investigations of the roles and interactions of genes, environment and plasticity for evolution.

**Supporting Information**

**S1 Fig**

**Relative expression of *nattl* and *nattl 1-3* in tissues of adult AC-charr.**

Relative expression of *Natterin* (A) & *Natterin paralogs 1-3* (B-D) within different tissues (skin, heart, liver, gill, spleen, intestine & kidney) of adult aquaculture charr (RT-qPCR); expression plotted for different tissues, relative to heart tissue (lowest expression levels).

**52 Fig**

**Relative expression of selected craniofacial candidate genes.** Relative expression of 12 candidate genes with characterized craniofacial expression during zebrafish development (ZFIN website) in the head of SB, LB, PL and AC at three time points in development. In the transcriptome data all of the genes had shown higher expression in SB at 200 τs. The expression is normalized to the geometric means of two craniofacial reference genes (*ACTB* and *IF5A1*). Expression is relative to a replicate of AC morph at 200 (*τs*), set to one. Error bars represent standard deviation calculated from two biological replicates and each biological replicate contains homogenate of six heads.

**S1 File**

**Parameters and multiple testing corrected p-values for expression analysis.**

The file is tab-delimited and the columns are;

- “Unigene.Description”: the annotation for that gene / paralog group.
- “NR.contigs”: number of contigs with this annotation.
- “logCPM”: count per million, log-scale
- “logFC.morph”: Mean fold change between the morphs, log-scale.
- “logFC.T163”, “logFC.T200”, “logFC.T433”: Mean fold change for each timepoints compared to timepoint 141, log-scale
- “FDR.morph”: P-value for morph difference, mulitple testing corrected
- “FDR.time”: P-value for time differences, mulitple testing corrected
- “Contigs”: SalmonDB id for the contigs with the specific annotation.

**S1 Table**

**qPCR primers**

**S2 Table**

**Mapping of Illumina reads to S.salar EST data.** Number of reads aligning to salmon reference for each sample

**S3 Table**

**ANOVA results** Nine genes were tested for differential expression between whole Small Benthic and Aquaculture charr embryos, at two developmental timepoints (161 and 200 rs).

**S4 Table**

**ANOVA for *Natterin.*** ANOVA for relative expression levels of *Natterin* and *Natterin Paralogs 1-3* in Arctic charr whole embryos (among SB, AC and PL morphs) and tissues from adult AC-charr

**S5 Table**

**Gene Ontology analyses of derived SNPs in SB-charr.**

**S6 Table**

**Predicted effect of SNP-candidates differing in frequency between charr morphs**

**S7 Table**

**Data on candidate polymorphisms, primers and melting temperatures.**

## Acknowledgments

This project was supported by The Icelandic Center for Research (grant number: 100204011) to SSS2, AP, ZOJ and BKK, The University of Iceland Research/Doctoral Fund to JG and KHK and University of Iceland research fund to AP, SSS2 and ZOJ. We would like to thank Baldur Kristjánsson for help with genomic region and alignments, and especially Droplaug N. Magnúsdóttir, Guðbjörg Þ. Örlygsdóttir, Steinunn Snorradóttir and Ólafur Þ Magnússon at deCODE Genetics for help with the Illumina sequencing.

## Author contributions

- Conceived and designed the study: JG, AP, ZOJ, SSS, SRF, VHM, EPA.
- Sampling, crosses and rearing was done by SSS, BKK, ZOJ, KHK, VHM, AP.
- RNA extraction and RNA sequencing, by SRF.
- Analyses of RNA sequencing data, JG, AP.
- qPCR work, EPA, SSS2, VHM.
- SNP analyses JG, AP.
- SNP confirmation IMJ, KHK, AP.
- Comparative genomic analysis AP.
- Writing: AP, JG, EPA, VHM, SSS.
- Analyses: JG, AP, EPA, SSS2.
- Gathered the data: ZOJ, SRF, EPA, IAJ, KHK, SSS2.

